# Spatial and Temporal Patterns of Public Transit Aerobiomes

**DOI:** 10.1101/2025.04.21.649744

**Authors:** Russell J. S. Orr, Ola Brynildsrud, Kari O. Bøifot, Jostein Gohli, Gunnar Skogan, Frank J. Kelly, Mark T. Hernandez, Klas Udekwu, Patrick K. H. Lee, Christopher E. Mason, Marius Dybwad

## Abstract

**Background:** Aerobiome diversity is extensive, however community structure at the species level remains elusive. Likewise, microbiomes of public transit systems are of public interest due to their importance for health, though few studies have focused on these ecosystems whilst utilising shotgun metagenomics. Aerosol studies have focused predominantly on individual cities, with limited between-city comparisons suggesting specific community structures. Longitudinal studies show aerobiome diversity as dynamic, fluctuating during seasonal and daily cycles, though an annual cycle remains to be considered. Further, a bacterial bias has limited fungal aerobiome studies, with few considering both fractions collectively. As such, the objective of this study was to examine spatial and temporal patterns in the species diversity of public transit aerobiomes, with an emphasis on bacteria and fungi.

**Results:** Air samples taken over a three-year period (2017-2019) from six global cities were subjected to shotgun metagenomic sequencing. Improved classification databases, notably for fungi, applying stringent parameters for trimming, exogenous contamination removal and classification yielded high species-level resolution. Microbial diversity varied substantially among cities, while human and environmental factors, recorded in parallel, were of secondary significance. Bacteria dominated the public transit aerobiome with increased presence in cities with higher population densities. All aerobiomes had complex compositions, consisting of hundreds to thousands of species. Annual variation had limited significance on the public transit aerobiome diversity and community structure.

**Conclusions:** Cities were the most important factor contributing to diversity and community structure, demonstrating specific bacterial and fungal signatures. Further, a correlation between geographical distance and the genetic signatures of aerobiomes is suggested. Bacteria are the most abundant constituent of public transit aerobiomes, though no single species is globally dominant, conversely indicating a large inter-city variation in community structure. The presence of a ubiquitous global species core is rejected, though an aerobiome sub-core is confirmed. For the first time, local public transit aerobiome cores are presented for each city and related to ecological niches. Further, the importance of a robust bioinformatics analysis pipeline to identify and remove exogenous contaminants for studying low biomass samples is highlighted. Lastly, a core and sub-core definition of contaminant aerobiome species with taxon tables, to facilitate future environmental studies, is presented.

## Background

The diversity of aerobiomes is extensive, encompassing bioaerosols of countless prokaryotic and eukaryotic species; predominantly bacterial, fungal and viral [1–7]. Despite bioaerosols receiving increased attention in recent years [3], largely during the SARS-CoV-2 pandemic [8–11], the community structure and diversity of aerobiomes, particularly at the species level, remains elusive. Microbiomes of public transit systems are of particular interest due to their importance for public health, with subways alone accounting for 190 million estimated daily travellers globally in 2019, a 20% increase from five years earlier [12–15].

Technological advancements in high-throughput sequencing (HTS) of environmental DNA (eDNA) has prompted an increase in studies utilising amplicon- and shotgun-metagenomics to profile subway surface microbiomes, principally associated with the Metagenomics and Metadesign of the Subways and Urban Biomes (MetaSUB) International Consortium [16–23]. Profiling community structure and species diversity of the public transit aerobiome, however, has been less prolific. The relatively low biomass that can be recovered from air, and the associated genomic processing have been major obstacles limiting aerobiome studies when compared to comparative works on soil, water and surfaces [16, 24, 25]. Recent developments in high-volume air samplers [3], eDNA isolation protocols from electret filters [3, 26–28], and improved HTS library preparation methods [3], have fuelled an increase in amplicon- and shotgun-metagenomic air studies [29–38]. However, the aerobiome of public transit systems, and in particular subways, remain understudied ecosystems.

Aerosol studies have focused predominantly on individual cities [29, 32, 35, 37, 38] making it difficult to infer possible spatial effects on airborne microbial community structure. Only a limited number of studies have compared public transit surface or aerobiomes between cities, with some results suggesting city-specific community structures [16, 33, 39–41]. However, the importance of geographical distance and population densities remains to be considered. Second, temporal studies in the outdoor environment show diversity and concentrations of the aerobiome as dynamic, fluctuating during seasonal and daily cycles [1, 42–44]. Though a universal pattern for bacterial and fungal diversity is lacking, results indicate ecosystem specificity, both positively and negatively correlated to factors including temperature, humidity and pollution [1, 2, 42–47]. Few studies have evaluated temporal effects on community structure in public transit aerobiomes, where results to date do not indicate any clear correlation of bacterial diversity with temperature and humidity [32, 35, 48]. Fungal air diversity studies are sparse, with the limited results available suggesting a negative correlation with temperature and humidity [34, 43]. As of writing, no shotgun metagenomics study has considered the effects of annual cycles on aerobiome diversity. Amplicon data is limited and inconclusive as to the stability of bacterial and fungal species richness between years [43, 49], with any public transit environment yet to be studied. Thirdly, a bias toward bacteria has limited fungal aerobiome studies [34, 36, 50]; spatial studies of fungi only suggest a potential to categorise geographically separated populations [1, 43, 51–55], whilst temporal studies indicate seasonal variation [34, 43, 48]. Presently, few studies consider the bacterial and fungal fraction collectively [1, 2, 56], and specifically for the public transit environment [33, 48]. Further, a major barrier for interpreting the role of fungi in aerobiomes has been database biases; studies building reference databases solely from the limited number of complete fungal genomes result in underrepresentation, with ensuing low read classification and insufficient resolution at the species level [37, 48, 57–59]. Fourthly, environmental factors linked to urbanisation, such as humidity and temperature, have been extensively studied and correlated to bacterial and fungal growth [32, 60–63]. Likewise, studies suggest that human factors, such as buildings and population densities, have shaped distinct microbial communities [64–66]. Though presently, the significance of these factors in shaping the microbiomes of public transit systems is largely unknown. Lastly, a “core” urban microbiome was recently introduced to investigate microbial species prevalence in the global urban environment [16]. The study, focusing on public transit surface microbiomes, defined a global “core” as species present in >97% of samples and demonstrated 31 taxa as globally distributed above this threshold [16]. Currently, this definition is yet to be applied to a substantial public transit aerobiome dataset, nor has it been applied to the city-level, where microbiomes are more homogenous [33, 38–41].

In general, microbiome studies utilising shotgun sequencing, and in particular those concentrating on bioaerosols, have been limited in their ability to resolve species-level diversity questions due to multiple factors [67]. In addition to the points already raised, and where adequate methodological description is provided, we find that aerosol sample pools have often been relatively small with questionable representations, hampering conclusions regarding community structure [35, 57, 68]. Further, sequencing depth has often been low, resulting in an undersampling of metagenomic diversity [58, 68]. Further, low sequencing depth has often influenced downstream stringency; firstly, with insufficient length and quality thresholds for read trimming impairing species-level classifications [37, 57, 69], and secondly, with relaxed read classification parameters that have allowed for inflated assignments with increased false-positives that result in an overestimation of species numbers [1, 2, 37, 48]. To reiterate, unrepresentative databases for read classification, predominantly with a bacterial bias, have limited resolution of species diversity [37, 48, 57, 58].

One of the main obstacles in inferring the true underlying community structure and species-level diversity of aerobiomes has been in mitigating contaminants pre-, peri- and post-sampling [70]. This is particularly important for low-biomass studies where exogenous contaminants can become overrepresented during amplification and blur results [71–73]. Sterile field practices are primary steps in reducing DNA contamination, although they do not eradicate it, and as such the importance of control samples is imperative for downstream revision of reads [26, 70, 73]. The importance of adequate controls is heightened post-sampling, where laboratory processing can contribute DNA contaminants from reagents [71, 74–77] and neighbouring samples [70, 74, 78]. However, the principal source of exogenous contamination post-sampling is human and human-associated DNA from the sample handler themself [71, 72, 79]. As such, exogenous DNA contributed during processing may constitute taxa homologous to that of the environment under study [16, 71, 80–82]. Though studies take mitigating steps during sampling and wet-lab procedures, lacking read data for negative controls results in shortcomings during dry-lab processes [70, 83]. Sufficient sequence depth of negative controls is critical to avoid undersampling and a subsequent partial removal of contaminant reads [33, 35, 57, 68, 71, 73, 84]. Lacking or partial contaminant read mitigation may result in an increased homogeneity between samples and therefore an overestimation of species abundance and prevalence [71–73]. Interestingly, public transit shotgun air metagenome studies suggest that microbiomes are dominated by a few species and primarily those of known human associations [33, 35, 37, 42, 48].

As such, the main objective of this paper is to examine species diversity in public transit aerobiomes and in particular spatial and temporal compositional patterns; a three-year annual time-series (2017-2019) and large-scale sampling scheme allows this to be studied for the first time, with an emphasis on bacterial and fungal species. In addressing the main objective, we address secondary research goals: First, evaluating the effects of environmental and human factors on microbiome structure, utilising metadata collected in unison with samples. Second, determining the presence of a public transit aerobiome “core” at global and local levels, focusing on species prevalence. Lastly, highlighting the importance of robust bioinformatic pipelines to identify and remove exogenous contaminants from low biomass samples for inference of underlying biological signals.

## Methods

### Air sampling

Air samples were taken annually over a three-year period (2017-2019) from public transit hubs of six cities, in total 750 air samples (Table 1 & Table S1). For 2018 a total of 261 samples were collected and sequenced from six cities (Denver, Hong Kong, London, New York, Oslo, and Stockholm) and 80 localities. For 2019 a total of 239 samples were collected and sequenced from five cities (excluding Stockholm) and 71 localities. Further, we reanalysed previously published data for 250 samples, collected and sequenced using the same protocols, from public transit hubs of the same six cities and 76 localities from 2017 [33]. Negative (22) and positive (5) control samples were utilized for downstream contamination removal.

**Table 1.**
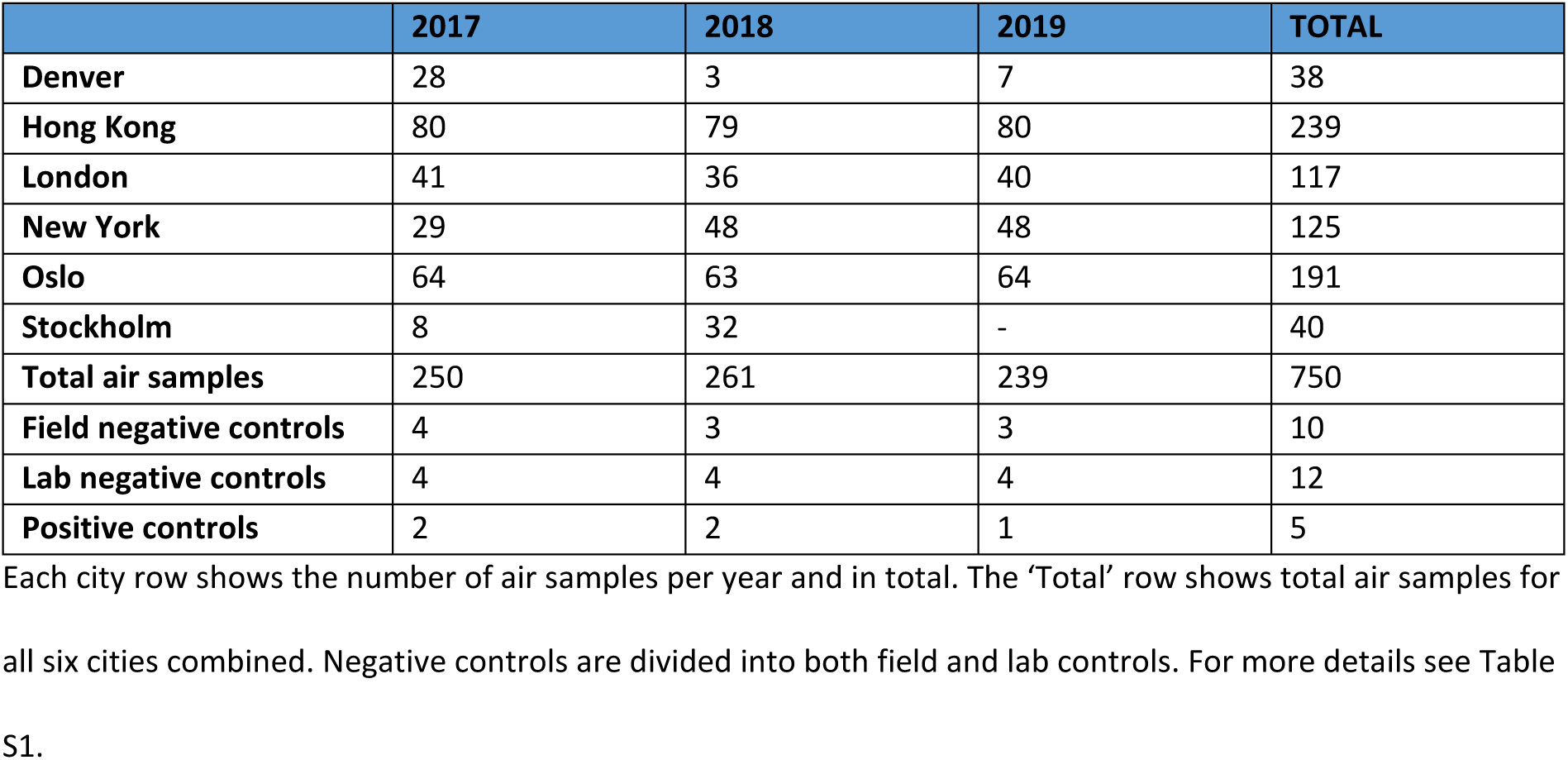
Number of samples per city per year.

Air sampling was performed yearly between June and August (Summer, Northern Hemisphere) at specific and repeated public transit localities in each city (Table S1). Samples were collected using a SASS3100 high volume electret filter air sampler (Research International, Monroe, WA, USA) with airflow set to 300 liters per minute for 30 minutes. The sampler was tripod mounted, facing 45° downward and 1.5 m above the floor. The aerobiome was collected on a sterilized electret filter (7100-134-232-01, Research International) and stored in a sterile 50 ml tube at −80°C. Further, metadata for a selection of sampling localities was recorded, including temperature (°C), relative humidity (%), number of travellers, whether the area was enclosed or open, and over- or underground (Table S1). To reduce contamination the SASS3100 was cleaned with alcohol wipes between samples. Gloves and disposable tweezers were used for filter handling. Allowing for characterisation and downstream processing of contaminants, “field” negatives (Table S1) were taken by mounting a filter to the inactive SASS3100 for a few seconds before storage, as outlined above. Air filters were shipped to the Norwegian Defence Research Establishment (FFI - Kjeller, Norway) on dry ice and stored at −80°C prior to DNA isolation.

### DNA isolation and quantification

DNA was isolated following the protocol of Bøifot et al., 2020 [26]. The protocol is adapted for aerobiome sampling on electret filters to achieve complete lysis and maintain community structure for Illumina sequencing. In brief, filters were extracted in NucliSENS lysis buffer (BioMérieux, Marcy-l’Étoile, France) before centrifugation (7000*g* for 30 min). The pellet was enzymatically lysed with MetaPolyzyme (SigmaAldrich, St. Louis, MO, USA) prior to 3 minutes bead beating in ZR Bashing Bead Tubes (Zymo Research, Irvine, CA, USA) with PowerSoil Bead Solution (Qiagen, Hilden, Germany) and Solution C1 (Qiagen) using a Mini-Beadbeater-8 (BioSpec Products, Bartlesville, OK, USA). Inhibitor removal was performed with the Qiagen solution C2 and C3. The pellet was subsequently combined with the original supernatant before isolating DNA with the NucliSENS Magnetic Extraction Reagents kit (BioMérieux) following the manufacturer’s protocol, with two modifications: the volume of silica beads was increased to 90 µl and the DNA-binding incubation time increased to 20 minutes. Isolated DNA was quantified with a Qubit Fluorometer 3.0 (Life Technologies, Carlsbad, CA, USA) using the dsDNA HS-assay before storage at −80°C. Negative “lab” controls were included during DNA isolation to adjust for exogenous contamination during analysis (Table S1). Positive controls were isolated utilizing the ZymoBIOMICS Microbial Community Standard (D6300, Zymo Research) to evaluate library preparation, sequencing bias on community structure, and determine classification and contamination thresholds.

### Library Prep and sequencing

Isolated DNA was shipped on dry ice and sequenced at HudsonAlpha genome sequencing centre (Huntsville, AL, USA). Libraries were prepped using an in-house protocol and sequenced on a NovaSeq 6000 to give 150 bp paired end reads.

Raw reads for 2017, 2018 and 2019 are available through the respective NCBI Bioprojects PRJNA561080, PRJNA1129830 and PRJNA1132165. Mean raw read depth for the samples from 2017, 2018 and 2019 were 9 440 986, 76 439 607 and 71 155 864, respectively. Corresponding numbers for the negative controls were 1 713 484, 77 150 860 and 106 546 906. (Table S1).

### Bioinformatics

#### Qualification, trimming and human read removal

Raw reads were quality controlled using FastQC v0.11.9 [85]. For high resolution species-level classification, reads were trimmed with Trimgalore v0.6.7 [86] with a length cut-off of 130 bp, a quality minimum of phred 30 and maintaining read pairs. Bowtie2 [87], in local mode, in unison with SAMtools v1.15 [88] removed reads mapping to the *Homo sapiens* (GRCh38) and phi X 174 genomes, again maintaining read pairs.

#### Sequenced diversity estimates

A redundancy-based K-mer approach to estimate average metagenomic coverage and sequence diversity per sample, excluding controls, was applied using Nonpareil 3 [89] with default parameters.

#### Cross kingdom and species level database construction and read classification

A Kraken2 [90] protein database, for cross-kingdom classification, was constructed from the entire NCBInr database. A Kraken2 and Bracken [91] nucleotide database (hereafter FBAV), for species level classification, was constructed utilising genome_updater v0.4.0 [92] to include Archaea, Bacteria and viral refseq complete genomes (871, 63 568 and 14 018 accessions respectively). In addition, reference and representative fungal genomes from Genbank and refseq assembled to complete genomes, chromosomes and scaffolds (3 049 accessions) were utilised. To gauge a complete eukaryotic and prokaryotic overview of community structure for each separate aerobiome sample, a cross-kingdom classification was performed; read pairs were classified to higher taxonomic levels (domain and kingdom) with Kraken2 utilizing the protein database (NCBInr) under default parameters. Classification to species level was performed with read pairs classified with Kraken2 utilizing the FBAV nucleotide database with confidence set to 0.1 and minimum-hit-groups to 4, to minimise false positives. Upon evaluation of the FBAV Kraken2 read classification reports for positive control samples, with known species structure, a 0.005% total read threshold cut-off was introduced to further minimise false positives [93] and applied using a modified Bracken script [94] to estimate species-level abundance.

#### Exogenous contaminant removal

Kraken2 classification reports for negative control samples were aggregated using an in-house script [95] to determine contaminating taxa for removal from air samples. Taxa defined as “contaminant” had to be present in at least two negative samples and have a total read count >10,000, representing a trade-off between sensitivity and specificity of detection. Contaminants present in lower abundance were deemed to have minimal impact on the results. Contaminating taxon ids (Table S2) were subsequently removed from read count tables with an in-house script [96]. For calculations on the core microbiome, taxa were only counted if their prevalence in air samples were significantly higher than their prevalence in negative control samples [73]. This was calculated with a two sample Z-test of proportion [97].

#### Read count normalization

To consider comparative species diversity metric calculations read counts were normalized to 10,000,000 per sample. Further, and to reduce the impact of highly abundant species on results, normalized reads were logarithmised. A pseudocount of 1 read was added to all taxa to avoid a logarithm of 0. Uniform Manifold Approximation and Projection (UMAP) [98, 99] was used for dimension reduction prior to visualization.

#### Aerobiome diversity

Species alpha-diversity of biological samples was calculated using the Shannon Diversity Index from the vegan package in R [100]. The Agricolae [101] package was used to run Tukey’s Honest Significant Difference test between groups, using city and year as group variables. Between-sample species (beta) diversity was calculated using UMAP, as previously described, and cross-contamination of samples between cities was evaluated from the result before exclusion and reanalysis (iterative).

#### Significance of city, year, environmental and human factors on community structure

The effect of city, year, temperature, relative humidity, traveller number, and whether the sampling point was located above or below ground (ground level; Table S1), on microbiome composition was investigated using Multivariate Analysis of Variance (MANOVA) [102].

Temperature was binarised to above/below 25 °C, relative humidity to above/below 65% and travellers to above/below 100. The top 10 components (total 32.1% of variance) from principal component analysis were used as dependent variables. Independent variables were chosen in a forward stepwise manner, using the F test to evaluate model improvements. F was calculated using Pillai’s trace. Variables contributing more to F were included before those that contributed less. Interaction terms with CITY were also included for all environmental variables. To be included in the final model, terms had to be significant at 0.001 threshold.

#### Relative classified read abundance

Kraken2/Bracken taxonomic read count tables against the FBAV database, minus contaminating taxa, and associated metadata per sample were imported into GRIMER v1.1.0 [71] for visualization. Relative classified read abundance was calculated per city, merging years and sampling sites, in addition to “total” for cities combined; firstly, at the kingdom/domain level to establish read abundance classified to Archaea, Bacteria, Fungus and Virus. Secondly, at the species level, showing reads classified to the top 20 species, in addition to those excluded as “other species”, for the complete microbiome (FBAV), before dividing to bacterial and fungal abundance separately.

#### The core public transit aerobiome

To determine presence of a “global core” public transit aerobiome, species prevalence for the complete microbiomes (FBAV), bacterial microbiomes, and fungal microbiomes were determined separately. The number of species for different prevalence definitions of “core” and “sub-core” [16] were evaluated under separate levels of species abundance; 0.005%, 0.3, 0.5% and 1%. To test possible effects of under sampling (low sequencing depth) on the global core, the analysis was rerun utilising only samples with an estimated average coverage >50% (i.e. > 50% of unique DNA in the sample estimated to be present in the filtered sequence file), established from the Nonpareil 3 result (2.4.2 and Fig. S1). Further, and considering more homogeneous environments, “local core” public transit aerobiomes (FBAV) were calculated for each city separately.

## Results

### Trimming and human read removal

Sequence reads removed from downstream analysis during primary quality trimming was on average 15.4% for air samples, with minimal deviation between cities. However, a lower mean read quality for 2017, and as such a larger read removal, with 38.0% of reads removed, was observed. In comparison, 13.7% of reads for 2018 and 14.3% of reads for 2019 were removed based on quality filtering (Table S1). For reads mapping to *H. sapiens* (GRCh38) and phi X 174 genomes, a mean of 24.0% (0.75%-94.6% range) were removed from downstream analysis for air samples. Variation between cities ranged from 14.3% for New York to 33.4% in Hong Kong. Negative control samples showed a mean total of 4.6% reads mapped to *H. sapiens* (GRCh38) and phi X 174 genomes (Table S1).

### Diversity estimates

Nonpareil estimates of metagenomic coverage and sequenced diversity (Fig. S1), post trimming and human read removal, showed that the 2018 and 2019 air samples, with comparative sequencing depth, had a similar estimated coverage. 2018 had a range of 0.27 – 1.00 with a mean of 0.52, whilst 2019 had a range of 0.23 - 0.99 with a mean of 0.50 (Table S1). The 2017 air samples, for comparison, had approximately 8x lower sequencing depth and an estimated coverage ranging from 0.13 to 0.46 with a mean of 0.25 (Table S1). Likewise, control samples for 2018 and 2019 had predominantly >0.99 sequenced diversity, with 2017 being undersampled at <0.80.

### Cross-Kingdom classification

An initial cross-kingdom metagenomic read classification, post trimming and human and phi X removal, to all eukaryotes and prokaryotes from the entire NCBInr protein database showed a conserved pattern between years (Fig. 1). The total result for all three sampling years combined (Fig. 1 & Table S3) shows 21.8% of reads as unclassified. The reads classified as prokaryotic, 64.9%, were divided into Bacteria (64.2%), Archaea (0.4%) and Viruses (0.2%). Eukaryotes comprised 13.3% of total classified reads, split into 7.7% Fungi, 2.5% Metazoa, 2.0% Plants and 1.1% as other eukaryotes. A deviation from the above result was 2017, with a moderately lower number of unclassified reads (20.1%), a high number of reads classified to bacteria (69.3%) and a lower number to Fungi (5.1%).

**Figure 1.**
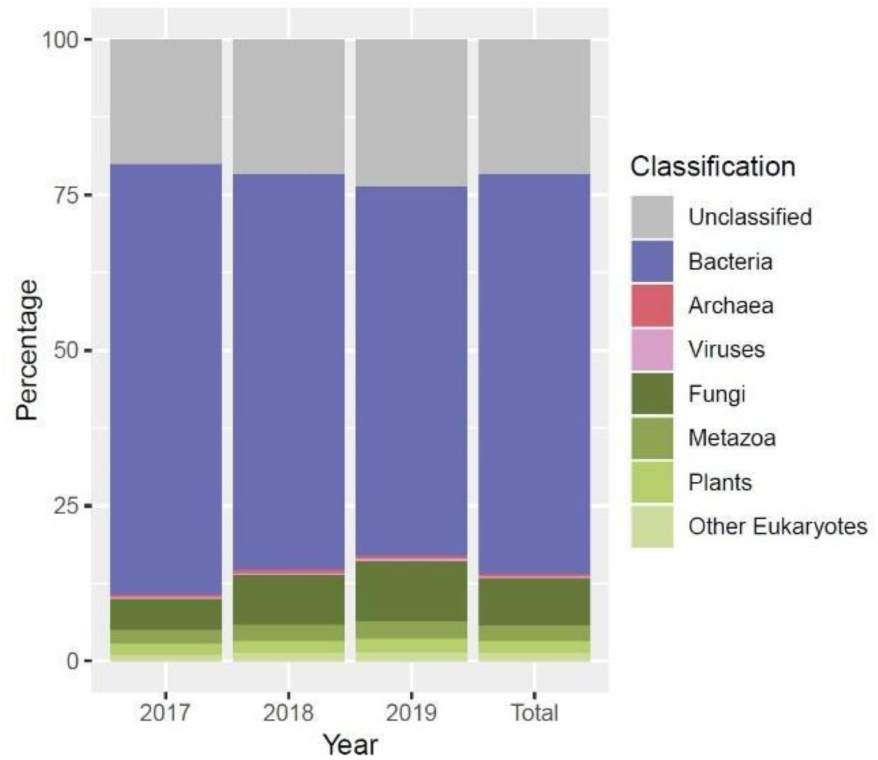
Higher level taxonomic classification to the NCBInr database: Metagenomic reads post trimming, and *Homo sapiens* and phi X removal, classified to the NCBInr protein database with default Kraken2 parameters. The chart shows year on year and total for higher taxonomic levels (domain and kingdom) for classified and normalised read counts: Archaea, Bacteria, Virus (prokaryotes) and Eukaryotes. The latter is further subdivided into Fungi, Metazoa, Plants and “other”. Unclassified reads are also depicted.

### Exogenous contaminant removal

Contaminating exogenous taxon ids were identified from combining all negative controls with the presented classification pipeline and removed from read count tables before further analysis (Table S2). Beta-diversity of control samples showed no statistically significant difference to warrant separation of field- and lab-negatives post trimming and human and phi X read removal (Fig. S2). Contaminants comprised a total of 290 taxa - 265 bacterial and 25 fungal species (Table S2 & Fig. S3). No contaminating Archaea or viral reads were identified. Contaminating taxa constituted a mean of 62.7% (55.6-70.7%) of the total classified reads from air samples (14.1-89.6%), of which 58.9% (51.0-66.3%) were identified as bacterial and 3.8% (1.6-6.7%) as fungal (Fig. 2A & Table S1). Accordingly, classified reads retained from the air samples for downstream analysis constituted a mean of 37.3%. For the 2018 and 2019 negative control samples 96.4-99.9% of reads were removed, with the remaining not classified as contaminants (Table S1 & Fig. S3). For the 2017 samples, with >50x lower negative control read coverage, this ranged from 76.5-96.4%. Conversely, positive controls for 2018 and 2019, which had higher DNA content, had a negligible read contamination of <0.01% (Table S1). For comparison, the 2017 positive control samples, with a >30x lower read coverage, had a mean read contamination of 0.4%. No correlation was observed between contaminant reads with city or year (Fig. 2A & Table S1).

**Figure 2.**
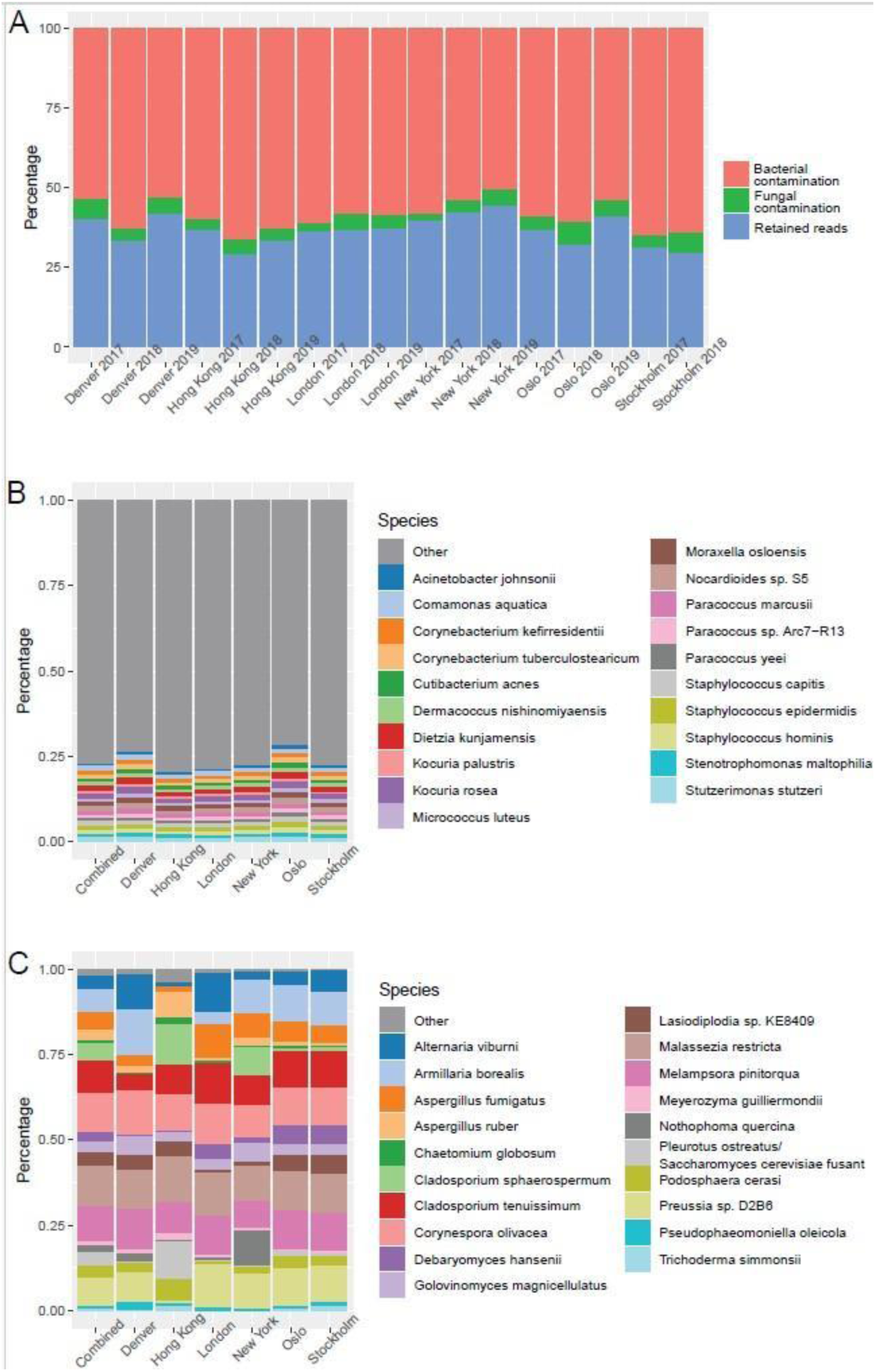
Air sample contamination and top contaminating species: A) Percentage of reads shown as bacterial and fungal contaminants, in addition to those retained for downstream analysis, for city and year. B) Relative abundance of the top 20 bacterial contaminants in the air samples in total and per city. C) Relative abundance of the top 20 fungal contaminants in the air samples in total and per city.

Relative abundance plots of the top 20 contaminating bacterial taxa for air samples illustrated an even distribution of species, with no single species dominating (Fig. 2B & Table S4); the most abundant taxon comprised only 1.4% of reads with the twentieth at 1.0%, whilst those outside the top 20 constituted 76.9% of total bacterial read contamination. The top three bacterial contaminants in the air samples identified by read number were *Micrococcus luteus*, *Cutibacterium acnes* and *Stenotrophomonas maltophilia*. Conversely, 98.5% of the relative read contamination for fungi was limited to the top 20 taxa, with a more uneven species distribution; The top five species (*Malassezia restricta, Corynespora olivacea, Melampsora pinitorqua, Cladosporium tenuissimum* and *Preussia* sp. D2B6) constituted >50% of read contamination (Fig. 2C & Table S5). Considering the prevalence of the identified contaminating taxa for all 750 samples (Table S6 & S7), four bacterial species were present in 100% of the samples: *C. acnes*, *M. luteus*, *Moraxella osloensis* and *S. maltophilia.* In >97% of the samples, previously defined as “core” [15], an additional 10 bacterial (*Acinetobacter johnsonii, Corynebacterium kefirresidentii, Corynebacterium tuberculostearicum, Dietzia kunjamensis, Kocuria rosea, Kocuria palustris, Paracoccus marcusii, Paracoccus sp, Arc7-R13, Staphylococcus epidermidis* and *Staphylococcus hominis*) and two fungal species (*Corynespora olivacea* and *M. restricta*) were observed, a total of 16 contaminants. Considering the >70% threshold, defined as “sub-core” [15], a total of 70 bacterial and five fungal contaminants were present prior to taxon removal.

Upon primary evaluation of beta diversity analysis (result not shown), atypical inter-city clustering identified 10 samples as possible cross-contaminants between cities. However, only a single 2018 sample from London (SL467228) that clustered with New York was deemed a true cross-contamination upon evaluation of metadata; the sample had unmeasurable DNA concentration and was located adjacent to New York samples on the library preparation plate (Table S1). It was subsequently excluded from downstream analysis. For the 9 remaining samples identified as possible cross-contaminants, no metadata (Table S1), e.g., lacking DNA concentration and sequencing plate position, supported an exclusion.

In total, 3 577 species were classified to the FBAV database from all public transit air samples in this study (Table 2 & Table S8). Of this, 2 560 were bacterial, 956 fungal, 36 viral and 25 Archaea species. Hong Kong had the highest number of total observed species (2 610), bacterial species (1 931), fungal species (653) and viral species (14). Though, Hong Kong also had the highest number of air samples (239). Conversely, Denver and Stockholm with the least number of air samples, 38 and 40 respectively, also had the lowest number of total, bacterial, fungal, viral and Archaea species (Table 2). A rarefaction analysis indicated a yet unsampled portion of species diversity in total and per city (Fig. S4).

**Table 2.** Number of classified species.

| City | Total | Bacteria | Fungi | Virus | Archaea |
| --- | --- | --- | --- | --- | --- |
| All | 3577 | 2560 | 956 | 36 | 25 |
| Denver | 980 | 776 | 199 | 2 | 3 |
| Hong Kong | 2610 | 1931 | 653 | 14 | 12 |
| London | 1132 | 836 | 276 | 4 | 16 |
| New York | 1673 | 1237 | 413 | 8 | 15 |
| Oslo | 1429 | 969 | 442 | 11 | 7 |
| Stockholm | 811 | 590 | 213 | 2 | 6 |

Each city and all cities in total (row), are divided to the studied kingdom/domain and total combined (column).

### Diversity estimates

The Shannon alpha diversity index shows species diversity (Fig. 3) based on the FBAV database. Tukey’s Honest Significant Differences test assessed within-city diversity variation per year and between-city variation. Within-city diversity appeared stable over the three-year sampling period, with no significantly different groups. However, we observe significant between-city differences, with New York having the highest species diversity and Denver the lowest. For the microbiome (Fig. 3A), London and New York were not significantly different, likewise, Denver and Oslo, whilst all other between-city comparisons were deemed significantly different. For bacteria and fungi separately (Fig. 3C & 3E, respectively), the following between-city comparisons were significantly different: Denver-Hong Kong, Denver-London, Denver-New York, Hong Kong-Oslo, London-Oslo, London-Stockholm, New York-Oslo, New York-Stockholm, Oslo-Stockholm.

**Figure 3.**
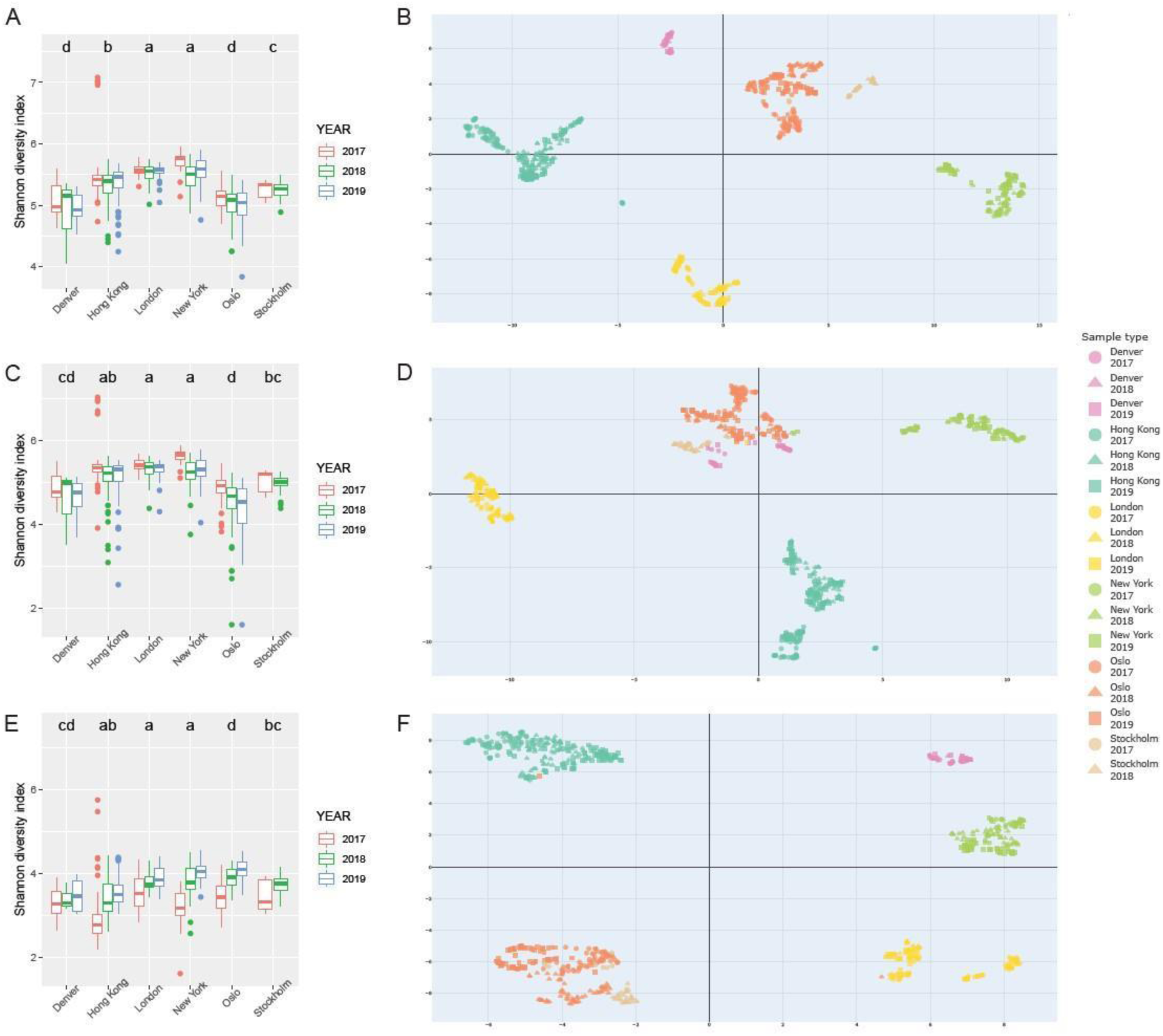
Alpha and beta diversity: Alpha (Shannon) diversity of the microbiome (A), bacteria (C) and fungi (E) is shown on the left; each city and year separate. Letters depicting Tukey’s Honest Significant Difference test between cities are shown at the top, where similar letters indicate no significant difference (p > 0.05). Beta diversity, with UMAP, for the microbiome (B), bacteria (D) and fungi (F) is shown on the right, each city and year separate. An interactive HTML version of each beta diversity plot (the microbiome (B), bacteria (D) and fungi (F)) is deposited as Fig. S5-7, respectively.

Beta diversity for the microbiome (Fig. 3B), showed clear clustering by city, and to a lesser extent a yearly clustering within cities. All six cities showed a distinct microbial composition signature, with only a handful of samples clustering discrepantly to their city of origin. For example, three Stockholm samples (SL342433-SL342435), clustered together with those from Oslo. To a lesser extent, six Hong Kong 2017 samples (SL310937-SL310942) form a cluster distant to that of the main city cluster. In all nine cases, multiple samples taken from the same localities on different days clustered within the main city grouping. As previous, no metadata (Table S1), e.g., lacking DNA concentration and sequencing plate position, supported an exclusion of these samples.

Considering bacteria separately (Fig. 3D), a strong city clustering was again demonstrated. Hong Kong, London and New York formed distinct groupings, albeit with the Hong Kong 2017 samples separating slightly from those of 2018 and 2019. Contrary to the complete microbiome result, we see Denver, Oslo and Stockholm forming a close but non-overlapping cluster for the bacteria-specific analysis. Within this cluster were two London samples (SL470243 and SL470310) and three New York samples (SL467183, SL470481 and SL470489). As previous, replicate samples from the same location group with the corresponding city cluster, and no metadata evidence supported cross contamination (Table S1). Interestingly, the three Stockholm samples (SL342433-SL342435) that clustered with Oslo in the microbiome plot (Fig. 3B) did not when considering bacteria. The six atypical Hong Kong samples (SL310937-SL310942), however, continued to form a cluster distant from that of their respective city cluster.

A similar pattern was shown for fungi (Fig. 3F), with a strong city community signal separating Denver, Hong Kong, London and New York into clearly defined clusters. Comparable to the microbiome and bacteria plot, the fungi plot shows the Stockholm 2017 samples cluster within Oslo, with those from 2018 being proximally isolated. Two Oslo samples deviate from the city signal, with SL469872 clustering within the Hong Kong grouping, and SL469711 placing proximal to London. Again, additional samples grouped with the main Oslo cluster and no metadata supported cross contamination (Table S1). For the three Stockholm samples (SL342433-SL342435) placed within Oslo for the microbiome result, a comparable result is shown for the fungi plot. The six Hong Kong samples (SL310937-SL310942), that placed distal to the main cluster for both the microbiome and bacterial plots, group within Hong Kong for the fungal plot.

### Significance of city, year, environmental and human factors on community structure

The significance of city signal on community structure for the microbiome, bacteria and fungi was confirmed with MANOVA. The final explanatory model (Table S9) showed the following factors as significantly contributing to the microbial composition, by order of descending Pillai value, interpreted as most to least significant: CITY, GROUNDLEVEL and YEAR (all p<1.0E-4). For bacteria, the significant contributors in order from most to least significant were CITY, YEAR, GROUNDLEVEL and HUMIDITY (all p<1.0E-4). For fungi, the significant contributors in order from most to least significant were: CITY (p<1.0E-4), YEAR (p<1.0E-4), GROUNDLEVEL (p<1.0E-4), CITY:GROUNDLEVEL (p<1.0E-4), HUMIDITY (p<1.0E-4), CITY:HUMIDITY (p<1.0E-4), and TEMPERATURE (p<1.0E-3).

### Taxonomic read classification

The relative read abundance plots for higher taxonomic levels (domain and kingdom), classified to the FBAV nucleotide database, showed the microbiomes of all six cities were dominated by bacteria, with a combined total of 74.4% (Fig. 4 and Table S10). Reads classified to fungi were also abundant, being 25.3% of the combined total, while Archaea and viruses were in minority, both with <1% of the total reads. Denver (36.6%) and Oslo (38.0%), had a higher percentage of fungal classified reads compared to the other four cities, a result in contrast to Hong Kong that showed a relatively high bacterial read classification (84.0%).

**Figure 4.**
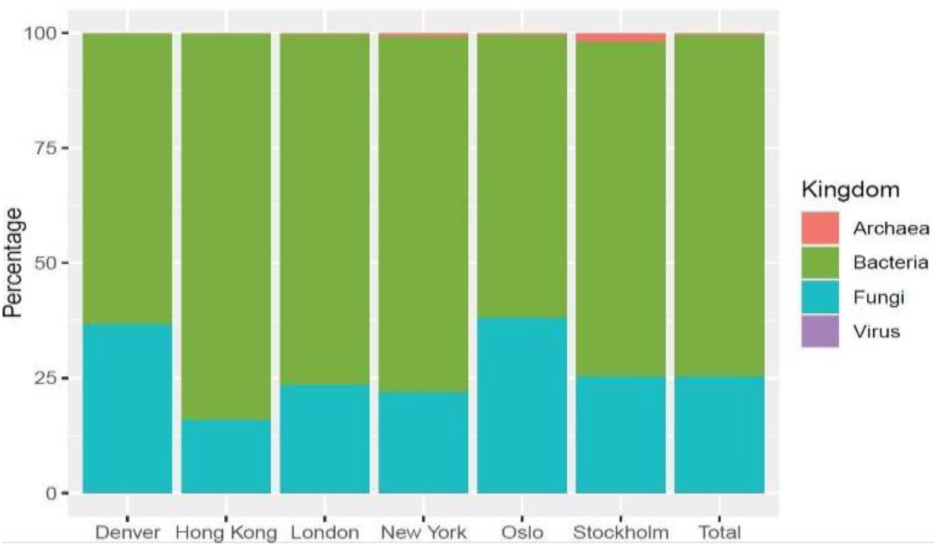
Relative classified kingdom/domain read abundance: For each city separately, and all cities in total for all sampling years combined. The chart shows the relative abundance of reads classified to Archaea, Bacteria, Fungi and Virus utilizing the FBAV database (Table S10).

Comparing relative read abundance at the species level for the microbiome (Fig. 5 & Table S11), showed 72.4% of diversity to be external to the 20 most abundant species, with large inter-city variation regarding taxa abundance. The top 20 species constitute 14 bacteria and six fungi, with Archaea and viral species unrepresented. No single species was shown as “globally” dominant, with the bacteria *Roseomonas mucosa* (3.5%) and the fungus *Ustilago bromivora* (2.2%) having the highest relative abundance. An increase in relative abundance for these species was seen when considering the cities separately, where *U. bromivora* had a higher abundance in Denver (24.7%) and London (5.5%), and *R. mucosa* was more abundant in Hong Kong (8.0%). Further, we observe relatively higher read classification to the bacterial species *Kocuria rhizophila* in London (6.8%) and *Nocardioides aquaticus* in Oslo (3.7%) and Stockholm (5.3%). Narrowing the read classification pool to solely bacterial or fungal species had no bearing on the result (Fig. S8 and Table S12 & S13).

**Figure 5.**
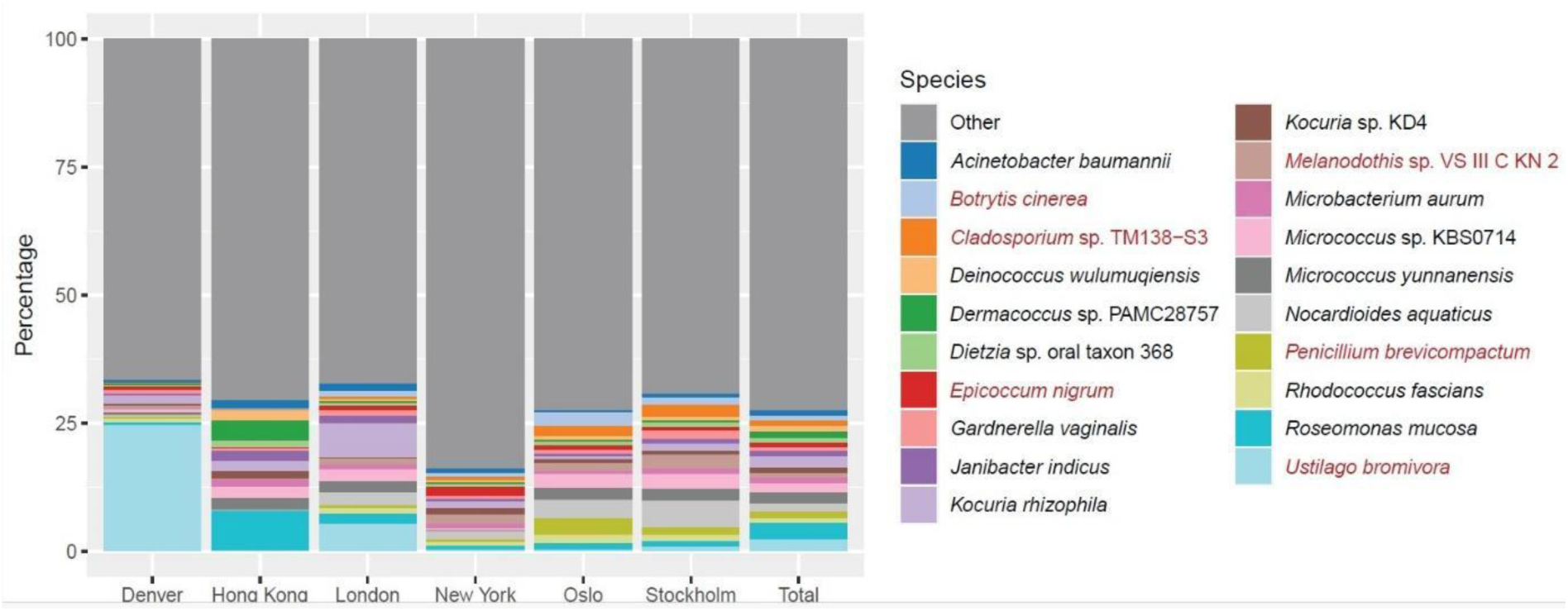
Relative classified species read abundance: The microbiome for all sampling years combined for each city and in total, classified to the FBAV database, with bacterial (black font) and fungi (red font) species. The chart shows the top 20 most prevalent species in addition to all “other” species (Table S11). The corresponding bacterial and fungal plots are provided as supplementary (Fig. S8 and Table S12 & S13).

### Core microbiome

#### Global core microbiome

To establish the presence of a “core” public transit aerobiome, the species prevalence of all air samples were considered (Fig. 6 & Table S14-21). No single species was present in >97% of air samples among the global pool (Fig. 6 & Table S14), a threshold previously defined as “core” [16]. The most prevalent species globally was *Dietzia* sp. oral taxon 368, present in 95.2% of samples. 44 species were present in the “sub-core” microbiome (Fig. 6 & Table S14), defined as 70-97% of samples [16]. The most prevalent species with high abundance (total abundance >1% averaged over samples) of the sub-core were *K. rhizophila* (94.8%), *Janibacter indicus* (86.3%), *Dermacoccus* sp. PAMC28757 (82.1%), *Acinetobacter baumannii* (78.3%), *Micrococcus yunnanensis* (77.9%), *Micrococcus* sp. KBS0714 (78.2%) and *R. mucosa* (76.6%). Bacteria dominated the public transit aerobiome sub-core, with only a single fungal species, *Cladosporiaceae* sp. IMV 00045, with a 72.3% sample prevalence and 0.9% abundance (Table S14). Many of the most prevalent species in the air samples were also present in the negative control samples, including the most prevalent species, *D.* sp. oral taxon 368. However, in all cases, the prevalence in the negative controls was lower than in the air samples, in contrast to the excluded contaminant taxa. To reiterate, homogenous species present in both the sampling and processing environment, e.g. air and human microbiome associates, were only excluded if an introduction post-sampling was statistically supported.

**Figure 6.**
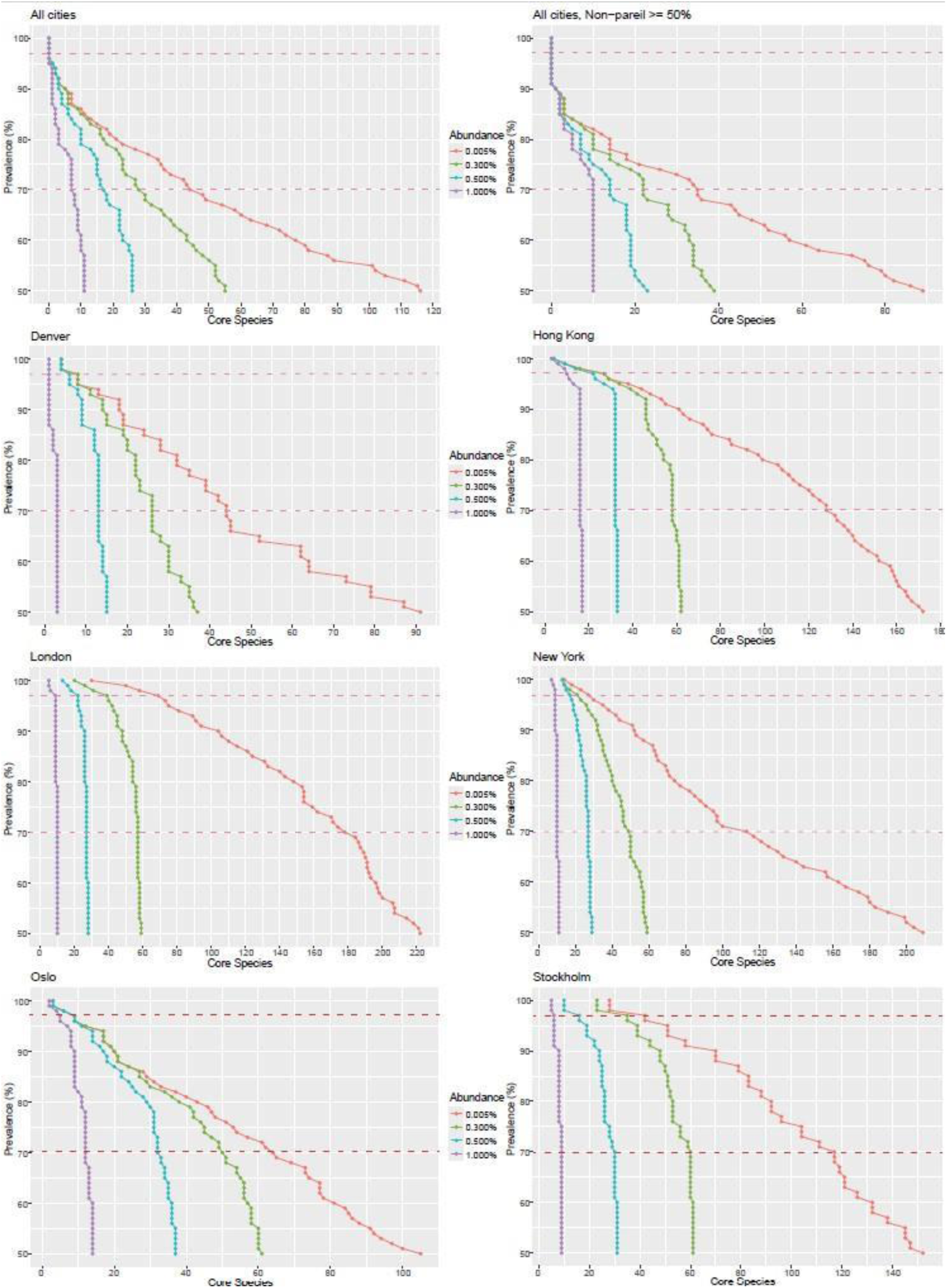
Core public transit aerobiome: The result shows the “global” core derived from all air samples (top left). The “global” core derived from air samples with an estimated Nonpareil diversity coverage >50% (top right). The bottom six plots show the core microbiome for each sampled city separately to consider effects of more homogeneous environments. The Y-axis shows % samples with species number on the X-axis, line colours depict species numbers at varying abundance levels from low (0.005%) to high (1%). Predefined prevalence definitions of “core” (97%) and “sub-core” (70%) are highlighted [16] with dotted lines. A complete list of “core” and “sub-core” species, prevalence and abundance levels for each inference is shown in Table S14-S21.

The global core microbiome analyses were replicated on a sample subset with a Nonpareil coverage >50% (231 air samples; Fig. 6 & Table S15) to study the effects of undersampling. For the Nonpareil high-coverage (>50%) sample set, congruent with the complete dataset, no species were present in the core, and again, the most prevalent species was *D.* sp. oral taxon 368, present in 90.5% of samples. For the sub-core, 35 species were recovered, where 12 species were lost and three gained compared to the complete dataset. Notably, *Brevibacterium* sp. CS2 and *Modestobacter marinus*, among the top 10 most prevalent species, were excluded from the sub-core, reducing from 90.0% to 67.1% and 86.5% to 62.0%, respectively. A general decrease in prevalence, and conversely increase in species abundance, was observed between the complete and reduced dataset.

#### Local core microbiome

The core microbiome analysis was replicated for each city separately to examine the presence of a “local core” aerobiome and the effect of more homogeneous environments on bacterial and fungal species prevalence and variance (Fig. 6 & Table S1).

The Denver core public transit aerobiome (Fig. 6 & Table S16) was dominated by the fungus *U. bromivora*, a grass pathogen, present in 100% of samples at an abundance of 15.8%. Additionally, in the core (>97%) we found *Clavibacter michiganensis* (100%), the pathogen *Clostridium botulinum* (100%), *Rhodococcus fascians* (100%), *Curtobacterium flaccumfaciens* (97.4%), *M. marinus* (97.4%), *Ornithinimicrobium flavum* (97.4%) and *Sphingomonas* sp. PAMC26645 (97.4%), though none were highly abundant (<1%). Denver’s sub-core microbiome constitutes 44 species.

The Hong Kong core microbiome (Fig. 6 & Table S17) was dominated by *R. mucosa*, present in all samples, with an average abundance of 7.31%. Other high-abundance species (>1%) in the core microbiome were *Dermacoccus* sp. PAMC28757 (100%), *K. rhizophila* (100%), *Deinococcus wulumuqiensis* (99.6%), *J. indicus* (99.2%), *D.* sp. oral taxon 368 (98.7%), *Kocuria* sp. KD4, (99.6%), *Microbacterium aurum* (98.7%), *A. baumannii* (97.5%) and *Nocardioides rotundus* (98.7%). Hong Kong was dominated by bacteria and only two fungal species were present in the core microbiome: *Penicillium sumatraense* (98.3%) and *Malassezia japonica* (97.5%), though both at <1% abundance. In total, the core microbiome for Hong Kong constituted 27 species and the sub-core 101.

The London core microbiome (Fig. 6 & Table S18) was governed by *K. rhizophila* (100%) with an abundance of 5.8%. Other high-abundance species in the core microbiome included *A. baumannii* (100%), *J. indicus* (100%), the fungal species *Melanodothis* sp. VS III C KN 2 (100%), *N. aquaticus* (100%), *Micrococcus* sp, KBS0714 (100%), *Aeromonas caviae* (97.4%), *M. yunnanensis* (97.4%) and *R. mucosa* (97.4%). In total, 69 species were included in the London’s core, of which four were fungal species, with the sub-core containing 109.

The New York core microbiome (Fig. 6 & Table S19) was again controlled by bacterial species, with the following present in 100% of samples at high abundance; *Arthrobacter* sp. QL17, *Deinococcus radiopugnans*, *Hymenobacter* sp. NBH84, *K.* sp. KD4, *M.* sp. VS III C KN 2, *Modestobacter* sp. L9-4 and *N. aquaticus*. However, the most abundant species of the core was the fungus *Epicoccum nigrum*, present in 99.2% of samples at 1.7% abundance. In total, the core encompassed 27 species, of which two were fungi, with the sub-core containing 86.

The bacterium *N. aquaticus* dominated the Oslo core microbiome (Fig. 6 & Table S20), comprising an average of 3.3% of all observations and present in all samples. However, and in contrast to other cities, Oslo had a high prevalence of fungi, with three highly abundant species present in the core; *Clavaria fumosa* (100%), *M.* sp. VS III C KN 2 (99.0%) and *Botrytis cinerea* (97.4%). The fifth and final abundant species was the bacterium *R. fascians* (99.0%). The remaining four species in the core were bacteria, and all in lower abundance, with the sub-core containing 55 species.

Congruent with Oslo, *N. aquaticus* (3.5%) was the most abundant species in the Stockholm core microbiome (Fig. 6 & Table S21), and likewise present in all samples. Additional core species of high abundance for Stockholm were the fungus *M.* sp. VS III C KN 2 (100.0%), and the bacterial species *Corynebacterium sanguinis* (100%), *Gardnerella vaginalis* (100%), *K. rhizophila* (100%) and *M.* sp. KBS0714 (97.5%). The total microbiome core for Stockholm consisted of 42 species, of which five were fungal, with the sub-core containing 75 species.

## Discussion

Air samples were taken annually over a three-year period during the summer months (2017-2019) from public transit hubs of six global cities: Denver, Hong Kong, London, New York, Oslo and Stockholm. Deep shotgun sequencing of air samples provides a substantial dataset for the study of spatial and temporal (annual) changes to the public transit aerobiome for the first time. As part of the study we built improved read classification databases, in particular for fungi, and applied stringent parameters for trimming, exogenous contamination identification and removal, and read classification to obtain high resolution at the species level. This high-quality dataset was used to study diversity at both the kingdom and species levels. We apply metadata taken in conjunction with sampling to infer the significance of city, year, environmental and human factors on community structure. Lastly, we evaluate the presence of a global and local core public transit aerobiome, with a focus on bacterial and fungal species.

### Diversity estimates

Annual summer sampling of the public transit aerobiome from six cities over three years shows a stable species diversity (alpha/Shannon); within city diversity is consistent across years for the complete microbiome (Fig. 3A) and was further validated when investigating bacteria (Fig. 3C) and fungi (Fig. 3E) separately. The aerobiome stability observed here, year on year, contrasts with previous studies on air and public transit aerobiomes investigating diurnal cycles and seasonal effects, which report significant temporal variation in bacterial and fungal diversity [1, 2, 32, 34, 35, 42, 44–48].

Inter-city spatial alpha diversity suggests a positive correlation between a city’s population density and the species richness of its’ aerobiomes (Fig. 3A); where cities with a higher population density, Hong Kong (6.3 K per km^2^), London (5.6 K per km^2^) and New York (11.3 K per km^2^), have richer microbiome communities than those cities with lower population densities, Denver (1.8 K per km^2^), Oslo (1.4 K per km^2^) and Stockholm (4.2 K per km^2^). Interestingly, bacteria (Fig. 3C) are contributing to this positive correlation between population density and species richness with fungal diversity comparatively constant (Fig. 3E). A result, comparable to that of soil microbiomes that also show a positive correlation between bacterial diversity and human population density [103]. The presented intra- and inter-city species diversity is congruent to that of results already published for the 2017 data, where 96.5% of reads were classified as bacteria [33]. A fungal comparison, however, is futile, as database limitations resulted in only 0.2% of reads being classified compared to 25.3% in the present study [33]. However, we show an inter-city result consistent with Abrego et al, 2020, who demonstrated with ITS barcoding that fungal species richness is not correlated with population density [55].

UMAP (Beta diversity) demonstrates a clear clustering to city (Fig. 3B), and to a lesser extent year, supporting the alpha diversity result and confirming that annual variation has a limited effect on species richness of the public transit aerobiome. The result indicates that underlying environmental aerobiome niches during summer months are relatively stable year on year when compared with seasonal variation [2, 32, 48, 49, 104]. Congruent with previous results, we infer a defined microbiome city signature, or fingerprint [16, 33, 39–41]. However, the separation between cities was more pronounced due to stringent methodological steps to remove contaminating reads and improve species classification.

For the microbiome (Fig. 3B), we can now postulate a relationship between geographical distance and genetic signature. For example, the cities Oslo and Stockholm, which have a close geographical proximity, have the most similar aerobiome populations. Considering the bacterial (Fig. 3D) and fungal (Fig. 3F) signal separately, the clustering pattern is more diffuse but congruent to that of the microbiome. For bacteria, Denver, Oslo and Stockholm form a close but non-overlapping cluster, which also lends support that cities with lower population densities have more homogenous and less diverse microbiomes than those of larger urban metropolises. Fungi, however, form clearly defined clusters with an overlapping community structure for Oslo and Stockholm, comparable with the microbiome. Currently, limited studies suggest the potential of city signatures for fungal aerosols; however, the potential to categorise geographically separated populations is supported by our result [43, 51–54].

Statistical significance of tested variables with MANOVA confirms city, congruent with beta diversity, as the most important factor contributing to microbial community structure. Year, as discussed, is less important but is still contributing slightly to the observed microbial composition. Further, though to a minor extent, GROUNDLEVEL, or if a sampling site is over- or underground, has a significant effect on community structure. GROUNDLEVEL is coupled with sites being either indoor or outdoor, which previous studies have shown to have distinct microbial communities shaped by factors including building design, ventilation and population density [64–66] Lastly, a significant correlation between bacterial and fungal composition to humidity is inferred. A positive correlation between humidity to bacterial and fungal growth has been extensively studied in addition to possible links to urbanization [60–63].

### Cross-kingdom taxonomic read classification

An initial read classification utilising the entire NCBInr protein database allowed for estimation of the total higher taxonomic diversity to all eukaryotes and prokaryotes. The public transit aerobiome appears stable over time with bacteria dominating (64.2%). Eukaryotic reads from fungi, metazoans, plants and other eukaryotes are less prolific, constituting 13.3% of the total relative classified diversity. Reads classifying to Archaea (0.4%) and Viruses (0.2%) are negligible, though the result is not adjusted for variance in genome size [105] and as such, is not an individual-count composition. Additionally, this study focuses solely on DNA and as such, possible presence of RNA virus in the public transit aerobiome is unaccounted for. Further, we observe a large unclassified diversity (21.8%), representing database limitations in addition to constraints of classifying genomic reads, which include a large non-coding portion, to protein databases. For comparison, though stressing differences in methodological approaches and protein databases, Gusareva et al., 2019, classified tropical aerobiome reads at the superkingdom-level to either bacteria (8%) or fungi (9%), with 83% unclassified [1]. Qin et al., 2020, studying Beijing air samples showed >93% of classified reads as bacteria and approximately 2% as eukaryotes [42]. Specifically for public transit aerobiomes, Zhang et al., 2024 showed >88% of classified reads as bacteria [35].

### Species-level taxonomic read classification

In this study, a total of 3 577 species were classified from 749 public transit air samples from six cities, with limited rare diversity yet to be sampled (Fig. S4). For comparison, Danko et al., 2021 classified a total of 4 424 species from 4 728 public transit surface samples from 60 cities [16]. For comparable public transit air studies; Zhang et al., 2024 classified 5 303 species from 18 samples from two locations in Shanghai [35]. Again, for Shanghai, Liu et al., 2024 classified 9 475 species from 108 samples and 4 stations [48]. Lastly, Lueng et al., 2021 analysing the same 250 samples from 2017 as this study, classified a total of 468 species-level taxa [33]. Though, methodological differences in sampling, sample processing and sequence analysis all negate direct comparisons between studies. As the above studies only report total species, a kingdom level comparison is lacking. For fungi, Abrego et al., 2020 report 976 species from 24 h air sampling of five Finnish cities [55], a result comparable to the 653 species identified in this study, which employed a shorter 30-min sampling period.

Removing exogenous contaminants and classifying nucleotide reads against a genomic clade specific database (FBAV), with increased stringency for species level classification, verified bacteria as most abundant (Fig. 4). However, we observe an increase in fungal classified reads compared to the cross-kingdom result, albeit to approximately 1/3 of that of bacteria, 25.3% against 74.4%. The relative increase in classified fungal reads may reflect nucleic databases being better suited for genomic read classification of eukaryotes, with a larger non-coding portion of the genome. Further, cities with lower population densities demonstrated an increase in relative fungal abundance, or conversely bacterial decrease, reflecting the Alpha (Shannon) diversity result (Fig. 1C), the result of Leung et al., 2021, in addition to reported results for urban soil microbiomes [33, 103]. A higher fungal abundance (25.3%) is demonstrated in this study compared to other public transit aerobiome studies that have classified reads to a nucleotide database; Liu et al., 2024, sampling the MTS in Shanghai had >95% of classified reads as bacterial, and approximately 4% as eukaryotic [48]. Similarly, Leung et al., 2021, analysing the same 2017 data as this study, had 96.5% of classified reads as bacteria, followed by virus (3.2%), fungi (0.2%), and Archaea (0.04%) [33]. This comparative discrepancy, however, is expected considering the larger database utilised in this study, resulting in an increased fungal, and conversely decreased non-fungal, classified proportion of the microbiome.

Interestingly, no single bacterial or fungal species was found to be statistically dominant in the public transit aerobiome when considering all six cities combined. Relative abundance between the top 20 most abundant species was relatively uniform and accounted for only 27.6% of the total public transit aerobiome diversity (Fig. 5), indicating that the public transit aerobiome is diverse. Our result contrasts with Leung et al., 2021 who infer two human associated species (*C. acnes* and *M. luteus*) as constituting approximately 50% of the public transit aerobiome when focusing on the same cities, though utilising only 2017 data [33]. As discussed, and in agreement with numerous low biomass microbiome studies [71, 72, 77, 79, 106], exogenous contaminants were excluded in this study, encompassing *C. acnes* and *M. luteus*.

Nonetheless, the human-associated bacteria *R. mucosa* [107, 108], is here identified as having the highest relative abundance for all air samples (3.5%), though it is also commonly isolated from environmental sources [109] and previously highlighted in comparable studies [33]. *R. mucosa* has a higher relative abundance (8.0%) in addition to sample prevalence (100%) when focusing purely on Hong Kong. The same species is present at 76.6% prevalence when considering all cities, and significantly lower for the negative control samples to be classified as a contaminant. The Hong Kong result, however, suggests a positive correlation between *R. mucosa* and the percentage of human DNA in the samples. Interestingly, this correlation is upheld when comparing cities separately, where those with lower levels of human DNA demonstrated reduced prevalence of *R. mucosa*. Inter-city variations in abundance are also observed for the prevalent bacterial species *K. rhizophila* in London (6.8%) and *N. aquaticus* in Oslo (3.7%) and Stockholm (5.3%), as previously highlighted by Leung et al., 2021 [33]. *K. rhizophila* lives predominantly in the rhizosphere, the soil encompassing plant roots, but it has a broad habitat, and highly adapted to ecological niches, being isolated from sources including mammalian skin, fermented foods, freshwater and marine sediments [110, 111]. Why *K. rhizophila* has a higher abundance in London, however, is surprising and may have a correlation with humidity, temperature and air change rate [112], though lacking metadata makes it difficult to conclude. *N. aquaticus,* however, is dominant in soil and wastewater and can endure low-nutrient conditions, and as such is known to degrade environmental pollutants [113]. A higher abundance of *N. aquaticus* in Oslo and Stockholm is likely associated with more optimal growth conditions including lower humidity and temperature [114] compared to Hong Kong, which had low abundance and prevalence of *N. aquaticus*.

*U. bromivora* was shown to be the dominating fungal species, with a 2.22% total abundance and 36.8% prevalence. Aerobiome studies from outdoor environments have identified *Ustilago* as abundant [2, 42, 55], while subway station samples have yet to identify this genus from either amplicon or shotgun sequencing data [32, 33, 37, 48]. This spore-forming biotrophic pathogenic fungus [115, 116] infects its plant host *Brachypodium distachyon*, a grass species with a broad native European, as well as invasive North American, distribution [117]. *U. bromivora* is abundant (24.7%) and prevalent (100%) in Denver, suggesting the presence of *Brachypodium grass* in the vicinity of the predominantly overground sampling sites. Surprisingly, a higher-than-average *U. bromivora* abundance (5.5% vs 2.2%) and prevalence (79.1% vs 36.8%) was also observed for London, a city with exclusively underground samples. Nonetheless, for all other cities the abundance of *U. bromivora* is <1%, with isolated increases inflating the total dominance of this species.

Similarly, we observe inflated abundance of the fungus *Cladosporium* sp. TM138-S3 in Oslo and Stockholm (2.0% and 2.6%), when compared to the overall result (0.9%). This global genus constitutes molds, which are found ubiquitously in the air and on decaying organic matter [118, 119]. The result may indicate a higher deterioration of building structures in the vicinity of the Oslo and Stockholm sampling sites [120]. To our knowledge, no other aerobiome shotgun sequencing study has considered fungal abundance at the species level whilst utilising a broad and representative database [1, 2, 33].

### Core microbiome

#### Global core microbiome

As species abundance only considers the total number of reads classified to a taxon, it may result in a biased and misleading interpretation as to which species are most broadly distributed, or prevalent. To consider if air sampled from global public transit hubs harbour a “core” microbiome, or community present in most samples, we instead focus on species prevalence. Previous work on a global public transit surface microbiome presented the definitions “core”; species present in >97% of samples, and “sub-core”; species present in 70-97% of samples [16]. The global public transit surface microbiome core constituted 31 bacterial species, while the sub-core comprised 1 145 species. Replicating the method on the public transit aerobiome, we find no support for a global core, with no single species present in >97% of samples. We do, however, confirm the presence of a sub-core, but this is limited to 43 bacterial and one fungal species.

Comparatively, Leung et al., 2021 found 17 species with prevalence >75%, however, of these only three are confirmed from this study (*Brachybacterium* with 57% prevalence, *Gardnerella vaginalis* at 84% and *K. rhizophila* at 95%), two (*Corynebacterium pseudogenitalium* and *Enhydrobacter aerosaccus*) are not recovered and may represent an incorrect classification, an additional five are flagged as contaminants (*C. tuberculostearicum, C. acnes, Cutibacterium granulosum, M. luteus* and *S. hominis*), and a further seven only have a genus-level classification (*Acinetobacter* sp., *Deinococcus* sp., *Dietzia* sp., *Kocuria* sp., *Massilia* sp., *Nocardioides* sp. and *Pseudomonas* sp.) that may or may not represent a contaminating species [33]. Congruent with the diversity result, species prevalence is dominated by bacteria, though, as fungi in general are more specialised to ecological niches than bacteria, a lacking global species prevalence may be expected [121, 122]. As such, a cosmopolitan generalist *Cladosporium* sp. is the sole fungal representative in the aerobiome sub-core [34, 43, 118, 119, 123].

The large discrepancy in results between public transit microbiome studies can be attributed to multiple factors; first, air and surface represent distinct microbiomes, where samples taken from the same sites have shown limited overlap, albeit for the most abundant and prevalent species [32, 124]. The relationship between air and surface microbiomes is complex, with biological and chemical properties, microorganism size, and environmental variables affecting both deposition and resuspension of microorganisms [32, 125]. Second, studies are not directly comparable, due to methodological differences pre-, peri-, and post-sampling impacting results [16, 70]. Third, surface swabs provide higher biomass than 30-minute air samples and are, therefore, subjected to fewer amplification cycles during preparation of sequencing libraries [32, 126]. As such, surface samples are less susceptible to over-amplification of exogenous contaminants, which remain negligible and require limited computation for identification and removal [16]. Highlighting the previous point, homologous taxa present as both constituents and contaminants of microbiome samples, predominantly species of human association, become even more complex to separate (see methods) with decreasing levels of biomass [71, 80–82].

We find that 24 of the 31 species in the surface core from Danko et al., 2021 have been excluded as exogenous contaminants in our dataset. This includes species discussed in detail later; the known human associates *C. acnes* and *M. luteus*, and the reagent contaminant *S. maltophilia*. Further, though Danko et al., 2021 utilised negative controls and highlighted both *Bradyrhizobium* sp. BTAi1 (also flagged in this study) and *C. acnes* as possible contaminants, they chose not to exclude them. Of the seven remaining surface core species (>97% prevalence), we reject the presence of *Pseudomonas stutzeri*, though this species has been reported previously [38]. The six other species were identified in the aerobiome: *Pseudomonas koreensis* (0.1%), *Sphingomonas taxi* (4.3%), *Brevundimonas* sp. GW460 (24.6%), *Lactococcus lactis* (67.7%), *J. indicus* (86.3%) and *K. rhizophila* (94.8%). However, only the two latter were included in the public transit aerobiome sub-core and represent the most prevalent species of the aerobiome, in addition to *D.* sp. oral taxon 368 (95.2%).

Our results suggest that, at a global level, no species are present at 100% prevalence in air. Danko et al., 2021 hypothesised that a 100% prevalence supported continued inclusion of the species highlighted above [16]. Though the only taxa present in our aerobiome dataset at 100% prevalence are four bacterial species, all of which were excluded as contaminants. Again, this includes *C. acnes*, *M. luteus* and *S. maltophilia*, in addition to *M. osloensis*, all of which are included in the surface core [16]. Further, a total of 14 bacterial and two fungal species that would otherwise have been included in the core public transit aerobiome were excluded as contaminants (Table S6 & S7). Additionally, 56 bacteria and three fungi when considering the sub-core were also excluded as contaminants. In comparison to non-aerosol studies [71, 127–130], that excluded seven bacterial species as contaminants (*A. baumannii*, *G. vaginalis*, *K. rhizophila*, *Prevotella copri*, *Prevotella melaninogenica*, *R. fascians* and *Schaalia odontolytica*), their inclusion as constituents of the public transit air sub-core is confirmed. As such, we propose an aerobiome contaminant core and sub-core (Fig. 2, Table S6 & S7), applying prevalence with the established definitions. Restricting the global analysis to high-coverage samples (Fig. 6), to test effects of undersampling, resulted in a decrease in the number of sub-core species but an increase in their abundance. This suggests that low-abundance samples inflate the number of species for low-prevalence definitions of the sub-core, highlighting the importance of sequencing depth and increased abundance cut-offs for interpreting prevalence.

#### Local core microbiome

As a global public transit aerobiome core is absent, we consider a local core, constituting the more homogenous environmental niches of each of the six sampled cities separately. For the first time, and in contrast to the global result, local aerobiome cores are present for each of the six cities. The local cores range from eight species in Denver to 69 in London and are dominated by bacteria, with fungal presence ranging from a single species in Denver to five in Stockholm. The uniqueness of the local core microbiomes reflects the diversity result (Fig. 3) and further highlights a cities location, or associated secondary components, as the most important factor contributing to microbial community structure [16, 33, 39–41]. The microbiome signatures of the geographically close Oslo and Stockholm are similar also when looking at the local core. The core microbiome, as with beta-diversity (Fig. 3), reflects how community composition depends on the ecology of the environment. Denver, for example, being surrounded by large areas of farmland has a core microbiome consisting of plant and soil-associated bacterial and fungal species. Notably, the pathogens *U. bromivora* and *Clostridium botulinum*. Likewise, Oslo and Stockholm, which have a higher fungus to bacteria proportion than the other cities, have a core of species associated with decaying plant and soil matter. On the other hand, Hong Kong, an urban metropolis, has a higher prevalence of bacterial and fungal species with a possible human association.

Though the data discounts the presence of a global public transit air core, and alternatively demonstrates local city cores, we highlight some limitations to the result; First, as the sampling period was both short and diurnal, it is plausible that species present in lower abundance for some sites were unrepresented. Second, possible effects of a sampling bias, e.g. differing sample numbers, degree of undersampling and unaccounted anthropogenic factors, and their impact on the core and sub-core cannot be disregarded. Though, adjusting for effects of undersampling may suggest otherwise, it also resulted in a significant reduction in the sample pool and species prevalence. Third, database biases impact classification, where reads may have been discarded as “unclassified” with a lacking species assignment. On the other hand, many earlier comparative studies have lacked a proper consideration of exogenous contamination and may have conversely overestimated both species’ abundance and prevalence [16, 33, 38, 48, 131].

### Contaminant filtering

#### Human read removal

When studying microbial communities with possible human association, the removal of *H. sapiens* reads prior to classification is fundamental. Sequence conservation between human, bacterial and viral genomes can result in partial mapping and incorrect interpretation of distribution and diversity [132]. Reads that mapped to the human GRCh38 and the phi X 174 bacteriophage genome assemblies were subsequently removed and accounted for a median of 23.8% for air samples and 6.3% for lab negatives. The result confirms that the dominant single source of DNA in the public transit aerobiome is of human origin, highlighting the importance of practices taken during sample processing [71–73, 77].

#### Exogenous contaminant removal

As amplification of low biomass samples result in over-expression of exogenous contaminants [71, 72, 77], screening and removal prior to downstream analysis is critical for recovering the true biological signal. To achieve this, reads or taxa present in the negative controls were removed from the air samples. One approach for exogenous contaminant removal is to first *de novo* assemble deep sequenced negative controls before excluding sample reads aligning to these contigs [33]. The success of this method, however, is directly correlated to the sequencing depth of the negative controls and their assembly quality; with low sequencing depth, or undersampling, resulting in the incomplete removal of contaminant reads from samples and subsequent misinterpretation of the data [33, 71, 73, 84]. In this study, we removed contaminating taxa from the air samples by adapting a method less sensitive to undersampling of negative controls [70, 71, 73]. Consequently, and supporting our methodological approach, negative control samples were stripped of >98% of their reads (Fig. S3 & Table S1). Conversely, positive control read counts and taxa distributions were negligibly affected. For the air samples, with lower biomass, a median of 64.7% of classified reads were removed in the decontamination step. Contaminating reads showed no significant difference between years or cities, again supporting an introduction post-sampling, but principally the importance of exclusion for inference of true biological signals.

In total, our exogenous contaminant removal pipeline identified and excluded 290 taxa - 265 bacterial and 25 fungal species, of which most are already flagged as common contaminants [71], including from other low biomass aerosol studies [1, 2]. The exclusion represents a total of 7.5% of the original species classification, 9.4% bacterial and 2.6% fungal, though as highlighted, predominantly taxa with high abundance and prevalence. Taxon ids of contaminants for future studies to utilise are provided (Table S2).

Contaminating taxa comprised known freshwater bacterial contaminants, including *Aquabacterium olei* and *Stenotrophomonas* spp. (in particular *S. maltophila* and *S. pavanii*), suggesting the “kitome”, with DNA present in extraction kits and reagents, as a possible source [71, 74–77]. Further, a potential “splashome” [74, 78], or sample cross-contamination, has been accounted for during the inter-city comparison conducted in this study. However, we predominantly observe known human-associated species such as *Corynebacterium* spp., *C. acnes*, *Dermacoccus* spp., *Micrococcus* spp. and *Staphylococcus* spp. (Fig. 2) [79, 133]. Likewise, fungal contaminants, though less abundant than bacterial, confirm human-associated species including *M. restricta* and *Meyerozyma guillermondi* [79, 134], in addition to common airborne molds such as *Aspergillus* spp. and *Cladosporium* spp. [135, 136]. We highlight the known species of the skin microbiome, *C. acnes* and *M. luteus* as examples emphasizing the importance of sequencing depth of negative controls, in addition to robust bioinformatic pipelines for the removal of exogenous contamination from low biomass studies. *C. acnes* and *M. luteus* are flagged as contaminants, introduced post-sampling, and subsequently excluded from downstream analyses. However, Leung *et al*., 2021, using the 2017 data consider both *C. acnes* and *M. luteus* as major constituents of the public transit air microbiome, representing approximately 50% of their reads [33]. Likewise, multiple aerobiome studies highlight the same species, in addition to other known and flagged human associates, as major constituents of aerobiomes despite minimal mitigation of contaminants pre-, peri- and post-sampling [35, 37, 42, 48].

A major challenge for low biomass studies, however, is to separate the true biological signal from that of an introduced contaminant [16]. This problem is heightened when both the environment being sampled in addition to the laboratory and reagents utilised post-sampling comprise homologous taxa, and in particular DNA from species of known human association [71, 80–82]. Nonetheless, when reads from human associated taxa are abundant in control samples, we presently find no other statistically viable method other than exclusion based on prevalence [71–73, 106], despite negligible numbers possibly representing a biological signal.

## Conclusions

This study contributes significantly to the advancement of air microbiomics and low biomass microbiome studies in general. In conclusion, cities were found to be the most important factor contributing to microbial diversity and community structure, demonstrating specific bacterial and fungal signatures. Further, a correlation between geographical distance and the genetic signatures of sampled public transit aerobiomes is demonstrated. Metadata was used to confirm that human and environmental factors (ground level and humidity) contribute significantly to the microbiome structure, however, to a lesser extent than that of cities. Bacteria are the most abundant constituent of the public transit aerobiome, with decreased presence in cities with lower population densities. No single bacterial or fungal species is globally dominant, conversely indicating the public transit aerobiome as rich and evenly distributed, with a large inter-city variation in community structure. No support is found, and as such the presence of a global core is rejected, though a sub-core dominated by bacteria is confirmed. For the first time, and in contrast to the global result, a local public transit aerobiome core is presented for each of the six cities and can be related to the available ecological niches. Further, the importance of a robust and extensive bioinformatics analysis pipeline to identify and remove exogenous contaminants when studying low biomass samples is highlighted and a contaminant core and sub-core definition and taxon tables for future studies to utilise is introduced. Lastly, annual variation is shown to have limited significance as to microbial diversity and community structure of the public transit aerobiome.

## Declarations

### Availability of data and materials

Raw sequence read sets for 2017, 2018 and 2019 are available through the respective NCBI Bioprojects PRJNA561080, PRJNA1129830 and PRJNA1132165. Supporting metadata is supplied in the supplementary tables 1-21.

### Authors’ contributions

**RJSO:** Lead, Study design, Data analysis, Data interpretation, Writing of manuscript (Lead writer). **OB:** Study design, Data analysis, Data interpretation, Writing of manuscript. **KOB:** Study design coordinated international sampling, Sample collection, Wet lab, Writing of manuscript. **JG:** Data analysis, Data interpretation, Writing of manuscript. **GS:** Coordinated international sampling, Sample Collection, Wet lab. **FJK:** Conception. **MTH:** Conception. **KU:** Conception. **PKHL:** Conception. **CEM:** Conception, Financing. **MD:** Conception, Study design, Financing, Writing of manuscript. **All authors**: Revision and approval of the final manuscript.

## Acknowledgements

**RJSO** thanks Ingjerd Thrane for assistance during wet lab procedures. Kristina Stenløkk for involvement in both sampling and wet lab protocols. **PKHL** thanks Marcus Leung and Xinzhao Tong for their assistance with the fieldwork in Hong Kong. **MTH** thanks Marina Nieto Caballero for assistance with sampling in Denver. **KU** thanks Nuno Rufino de Sousa and Antonio Rothfuchswith sampling in Stockholm.

## Supplementary Figures

**Supplementary Figure 1.**
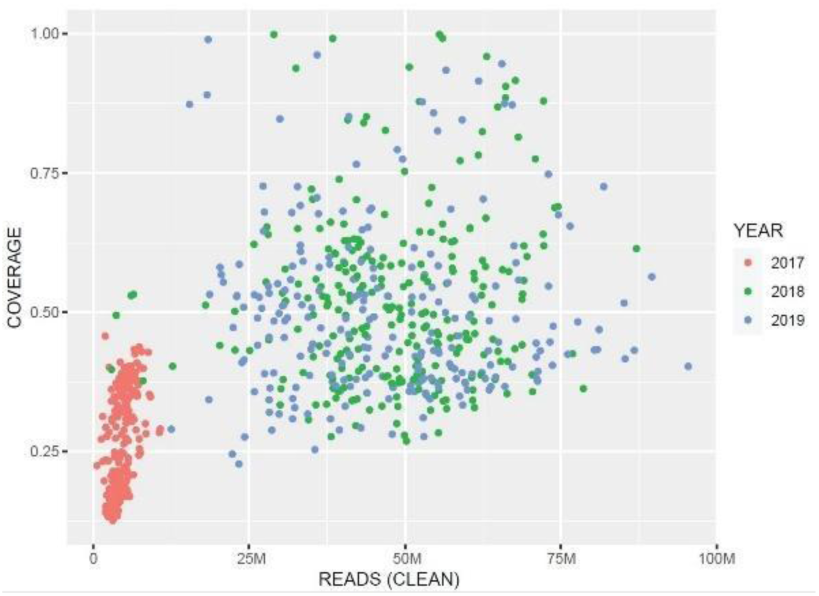
Nonpareil estimates of metagenomic coverage and sequenced diversity: post trimming and removal of *Homo sapiens* and phi X 174. Red dots depict air samples from 2017, green from 2018, and blue from 2019. Read number, in millions, is shown on the x-axis, with coverage on the y-axis.

**Supplementary Figure 2.**
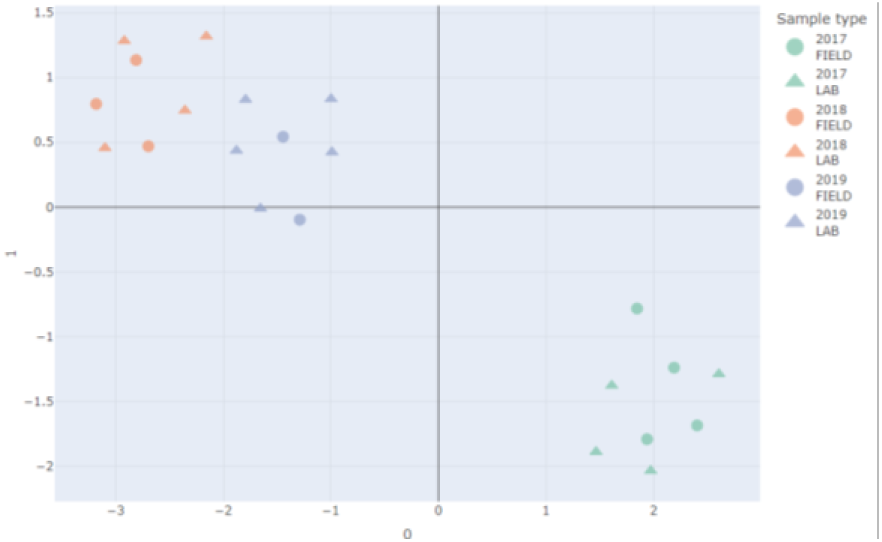
Beta diversity of negative control samples: Difference test with UMAP between field- and lab-negatives post removal of *Homo sapiens* and Phix reads and separated to year.

**Supplementary Figure 3.**
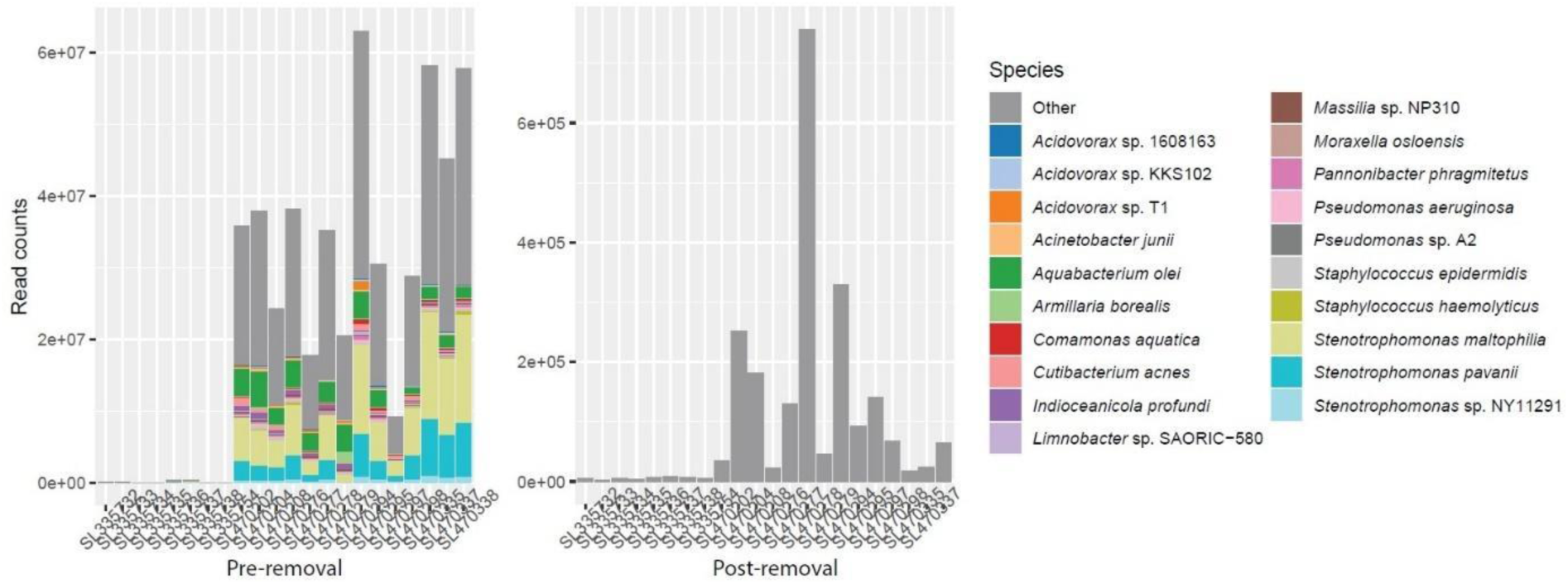
Relative read abundance of negative control samples pre- and post-contaminant removal. The top 20 microbiome contaminants annotated with colour for all negative samples pre- (left) and post-removal of contaminating species taxon IDs.

**Supplementary Figure 4.**
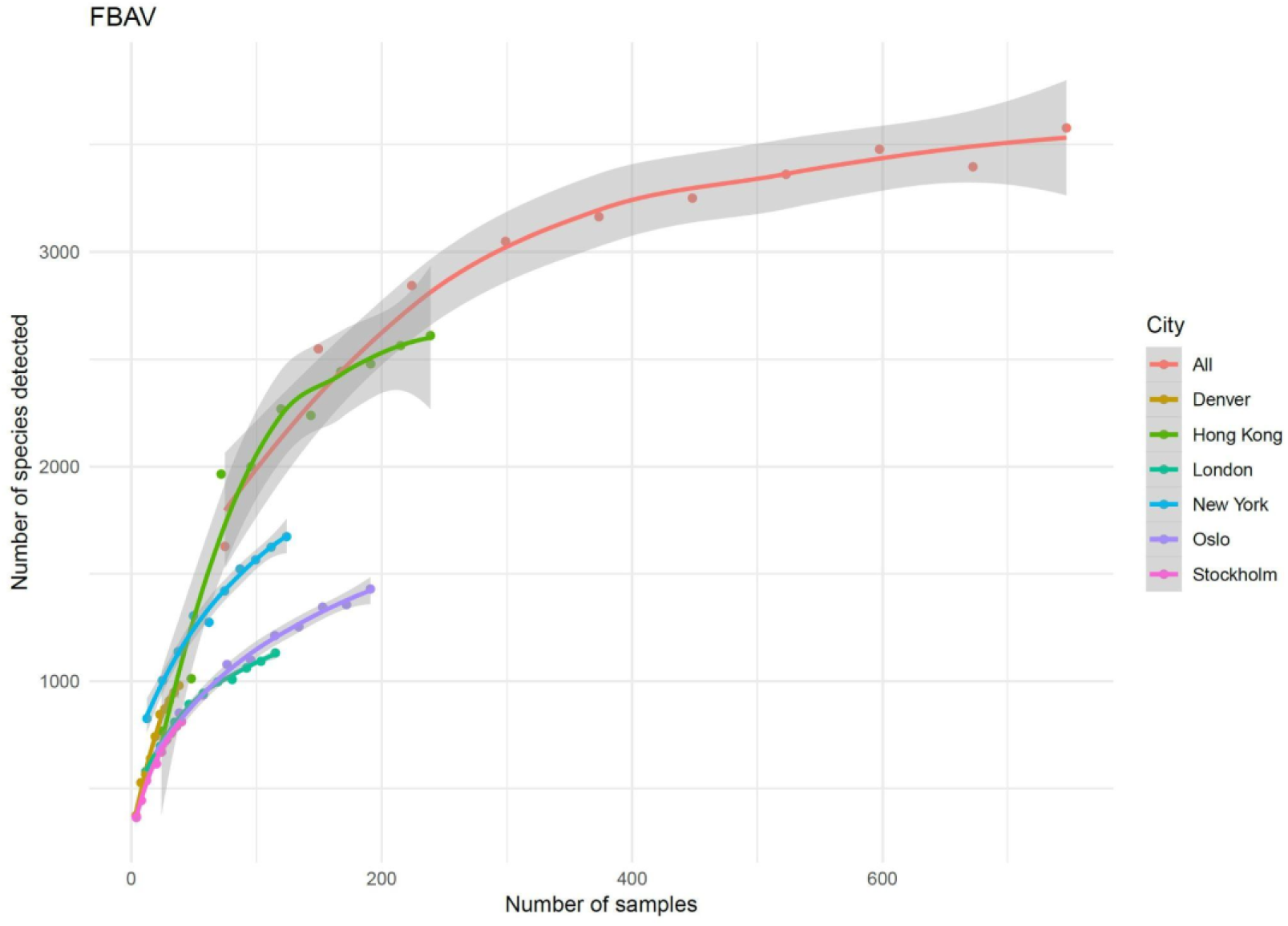
Rarefaction analysis of total sampled species diversity: For all cities combined and separately against the FBAV database, as assessed by subsampling each dataset, 1,000 permutations

**Supplementary Figures 5-7. Interactive HTML beta-diversity plots:** with UMAP, for the microbiome (Fig. S5), bacteria (Fig. S6) and fungi (Fig. S7) separated by city and year. Taken from Fig. 3

**Supplementary Figure 8.**
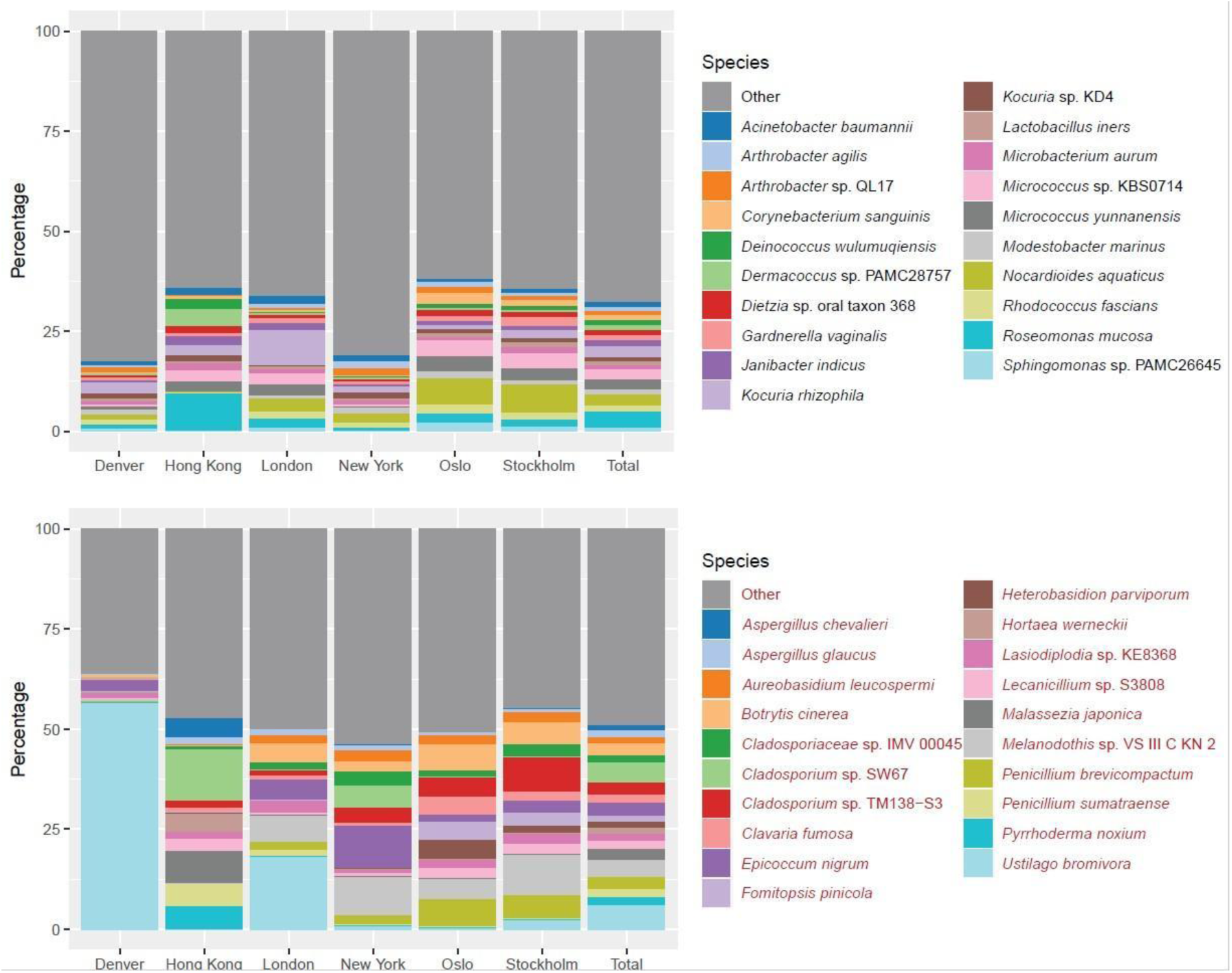
Relative classified species read abundance: To the bacterial (top) and fungal (bottom) microbiome for all sampling years combined, classified to the FBAV database. The chart shows the top 20 most prevalent species in addition to all “other” species. Charts are divided into cities and the total of all six cities combined.

